# Use of Mucosally Administered Outer Membrane Vesicles Derived from *Bordetella pertussis* to Diminish Nasal Bacterial Colonization

**DOI:** 10.1101/2024.03.11.584448

**Authors:** E. Rudi, E. Gaillard, D Bottero, D Hozbor

**Author notes:** Corresponding Author: Dr. Daniela Hozbor.

## Abstract

**Background:** We previously identified *Bordetella pertussis*-derived outer membrane vesicles (OMVs) as a promising immunogen for improving pertussis vaccines. In this study, we evaluated the efficacy of our vaccine prototype in immunization strategies aimed at reducing disease transmission by targeting colonization in the upper airways while maintaining protection against severe disease by reducing colonization in the lower respiratory tract.

**Methods:** We assessed different mucosal administration strategies in a murine model, including homologous 2-dose schedules and heterologous prime-boost strategies combining intramuscular (IM) systemic immunization with mucosal routes (intranasal (IN) or sublingual (SL)). We utilized mucosal c-di-AMP and/or systemic alum adjuvants to formulate the OMV vaccine prototype. A homologous IM immunization schedule and commercial vaccines were used for comparison.

**Results:** All tested heterologous schemes induced higher levels of specific IgG with significant avidity, as well as high levels of IgG1 and IgG2 compared, to corresponding homologous 2-dose schemes via mucosal routes (OMV_IN-IN_ or OMV_SL-SL_). High IgA levels were observed post-*B. pertussis* challenge following OMV_IN-IN_ treatments and heterologous treatments where the second dose was administered via a mucosal route. Furthermore, schemes involving the intranasal route, whether in a homologous or heterologous scheme, induced the highest levels of IL-17 and IFN-γ. Accordingly, these schemes showed superior efficacy against nasal colonization than the commercial vaccines. Specifically, homologous intranasal immunization exhibited the highest protective capacity against nasal colonization while maintaining an excellent level of protection in the lower respiratory tract. To enhance the protective capacity against nasal colonization further, we conducted a comparative analysis of formulations containing different adjuvants (c-di-AMP, alum, or a combination of both) administered via homologous intranasal routes. These assays revealed that the use of alum, either alone or in combination with c-di-AMP, did not enhance the immune protective capacity.

**Conclusions:** All the experiments presented here highlight that the use of OMVs, regardless of the scheme utilized, with the exception of OMV_SL-SL_, outperformed acellular pertussis (aP) vaccines, achieving a greater reduction in bacterial colonization in the upper respiratory tract (p<0.001).

## Background

Pertussis is a respiratory infectious disease that affects individuals of any age but is more severe in young children who are unvaccinated or have incomplete vaccination schedules. In unimmunized infants, the disease can be lethal. Characterized by bouts of violent coughing, vomiting after coughing, cyanosis, and apneas, the disease is caused by the Gram-negative bacterium *Bordetella pertussis* (1). Studies in animal models and even in humans have shown that a robust humoral immune response and IFN-γ, produced by Th1 cells, play a critical role in protection against the symptoms caused by primary infection with *B. pertussis* and also in adaptive immunity against reinfection (2–5). Recent studies using the non-human primate model (baboons) have shown that Th17 cells and IgA also play a role in protective immunity against this bacterium (6,7).

Vaccination is the preferred prevention strategy for this contagious disease. Currently, two types of pertussis vaccines exist, with the first one developed consisting of non-replicating cells from the causative agent (wP). This formulation that induces a potent Th1 and Th17 cell responses, as well as the establishment of a tissue-resident memory population (TRM) (8) is effective in preventing pertussis in children up to 7 years old. Adverse effects associated with wP vaccines led to their non-recommendation for the adolescent and adult population and prompted the development of a new generation of more tolerable vaccines based on purified *B. pertussis* immunogens (acellular vaccines, aP) (9–12). The aP vaccines were introduced in the late 90s into routine immunization programs in many developed countries. In recent years, the incidence of pertussis has increased in several countries, including those with high vaccination coverage (13–16). Various explanations have been proposed for the resurgence of pertussis, including improvements in integrated disease surveillance, the prevalence of circulating *B. pertussis* strains more resistant to immunity conferred by vaccination (especially that induced by acellular vaccines), and the loss of vaccine-induced immunity (which occurs more rapidly in the case of aP) (17–19). The resurgence of the disease has prompted a short to medium-term response, leading to the incorporation of additional vaccine boosters in the adolescent and adult population, particularly in pregnant women (20,21). The data collected so far indicate that pregnant women vaccination strategy is proving successful (22,23). However, challenges persist in vaccination strategies that require resolution. One such challenge is that current vaccines, particularly acellular vaccines, are incapable of reducing the risk of bacterial colonization in the upper respiratory tract (24,25). Consequently, disease transmission may still occur even in vaccinated individuals. Of greater concern is that vaccinated individuals who become infected and remain asymptomatic could silently transmit the disease (26–28).

Recently, it has been demonstrated that replacing alum with adjuvants promoting Th1 cells and/or TRM, including Toll-like receptor (TLR) agonists, can enhance the protective efficacy of experimental aP vaccines in mice (29–31). In particular, the combination of LP1569 (TLR2) and the agonist for the intracellular receptor stimulating interferon gene, c-di-GMP, in an aP formulation induced significant numbers of respiratory IL-17-secreting CD4 TRM cells and prevented nasal colonization for at least 10 months after immunization (31). In preclinical assays we have previously demonstrated that outer membrane vesicles (OMVs) derived from *B. pertussis* have the potential to be tested as a third generation of pertussis vaccines. They have proven to be safe, induce a Th1/Th17/Th2 profile, generate tissue-resident memory cells, and exhibit significant capability to reduce bacterial colonization in the lower airways (associated with severe disease). We detected this protective capacity with the OMV administered both systemically and mucosally (32,33). Raeven *et al* (34) conducted comparative studies between vaccination schemes using OMVs derived from another strain of *B. pertussis* administered subcutaneously and intranasally. The results obtained showed that intranasal administration of OMVs is an appropriate route to reduce transmission (bacterial colonization in upper tract) and severe disease (bacterial colonization in lower respiratory tract) when animal were exposed to bacterial suspension containing 10^5^ CFU (34). In this study, aimed at further establishing OMVs as a viable immunization alternative, we assessed the impact of administering OMVs via different mucosal routes, sublingual and intranasal, in both homologous (same route for different doses) and heterologous (different routes for different doses) schemes on immunogenicity and protective capacity against bacterial colonization in the upper and lower respiratory tracts. OMVs formulations with different adjuvants were also assayed. For comparison purposes OMVs, aP y wP administered by intramuscular route were also evaluated.

## Materials and Methods

### Mice

C57bl/6 mice (4 weeks old), obtained from the Faculty of Veterinary Sciences, La Plata, Argentina, were kept in ventilated cages and housed under standardized conditions with regulated daylight, humidity, and temperature. The animals received food and water ad libitum. The animal experiments were authorized by the Ethical Committee for Animal Experiments of the Faculty of Science at La Plata National University (approval number 004-06-15, 003-06-15 extended its validity until August 10, 2027).

### *B. pertussis* strain and growth conditions

*B. pertussis* Tohama phase I strain CIP 8132 was used throughout this study as the strain for challenge in the murine model of protection. Bacteria were grown in Bordet–Gengou agar supplemented with 10% (v/v) defibrinated sheep blood (BG-blood agar) for 72h at 36.5 ºC. Isolated colonies were replated in the same medium for 24h and then resuspended in phosphate-buffered saline (PBS: 123 mM NaCl, 22.2 mM Na_2_HPO_4_, 5.6 mM KH_2_PO_4_ in MilliQ® nanopure water; pH 7.4). The optical density at 650 nm was measured and serial 10-fold dilutions plated onto BG-blood agar to determine the number of bacteria in the challenge inoculum.

### Isolation and characterization of outer membrane vesicles (OMVs)

OMVs were isolated and characterized as previously described (32,35). Briefly, culture samples from the decelerating growth phase were centrifuged and the bacterial pellet obtained was resuspended in 20mM Tris–HCl, 2mM EDTA pH 8.5. The suspension was sonicated in cool water for 20 min. After two centrifugations at 10,000×g for 20 min at 4 °C, the supernatant was pelleted at 100,000×g for 2 h at 4 °C. This pellet was re-suspended in Tris buffer (20 mM pH 7.6). The samples obtained were negatively stained for electron microscope examination. Protein content was estimated by the Bradford method using bovine serum albumin as standard (36). The presence of the main immunogenic proteins in the OMVs was corroborated by immunoblot assays using specific antibodies as we previously described (not shown) (32,37).

### Formulation of OMV-based vaccine

The characterized OMVs that range in size from approximately 50 to 200 nanometers in diameter were used to formulate the vaccine with tetanus (5 to 7 Lf/ dose with a power greater than or equal to 2 UIA/ml serum) and diphtheria (1 to 3 Lf / dose with an output of 0.1 UIA/ml serum) toxoids as we previously described. OMVs formulated with alum (Alhydrogel 2%, CRODA), c-di-AMP (7.5 µg per dose) (Invivogen), or c-di-AMP (7.5 µg) combined with alum (Alhydrogel 2%) as adjuvants were evaluated throughout this study. To perform the experiments described below we verified that the OMV based vaccine prepared by us fulfilled the WHO criteria for safety in the weight-gain test https://cdn.who.int/media/docs/default-source/biologicals/vaccine-standardization/pertussis/annex-6-whole-cell-pertussis.pdf?sfvrsn=f734b4_3&download=true). The safety of OMV-based vaccines was also confirmed by human whole-blood assays (38).

### Immunization of mice

Groups of 4-8 female C57BL/6 mice were immunized with OMV-based vaccine formulated as previously described with 3-6 µg total protein per dose formulated with alum, c-di-AMP or a combination of both as adjuvant (discriminated in the legends to the figures), or 1:10 human dose of aP vaccine BOOSTRIX® [GlaxoSmithKline, with composition per human dose: pertussis toxoid (8 µg), pertactin (2.5 µg), filamentous hemagglutinin (8 µg), tetanus toxoid (20 IU), diphtheria toxoid (2 IU) and alum as adjuvant] or 1:10 human dose of whole cell vaccine (Serum Institute of India PVT LTD, composition per human dose is: Diphtheria Toxoid ≤ 25 Lf (≥ 30 IU), Tetanus Toxoid ≥ 5 Lf (≥ 40 IU), *B. pertussis* ≤ 16 OU (≥ 4 IU) adsorbed on aluminum phosphate ≤ 1.25 mg) or) using a 2-dose schedule. Two weeks after the last immunization blood samples were obtained. For protection assays mice were challenged with *B. pertussis* by intranasal inoculation (sublethal dose 1×10^7^- 5×10^7^ CFU 40μl^-1^) as is described below. Mice were sacrificed 1 week after challenge. Bacterial counts were performed 7 days after the challenge as described previously.

### Enzyme-linked immunosorbent assay (ELISA)

As we previously described (39), plates (Nunc A/S, Roskilde, Denmark) were coated with proteins from bacterial lysates at 3 µg/ml in 0.5 M carbonate buffer pH 9.5, by means of an overnight incubation at 4 ºC. Blocked plates with 3% milk in PBS (2h 37ºC) were incubated with serially diluted samples of mouse serum (1h 37ºC). Sera from different period were obtained after leaving the blood samples to clot for 1h at 37ºC followed by centrifuging for 10min at 6,000xg. IgGs from individual serum or pooled sera bound to the plates were detected after a 2-h incubation with goat anti– mouse-IgG–linked horseradish peroxidase (1:8,000 Invitrogen, USA). For measuring IgG isotypes, detection of bound antibody was determined using HRP labeled subclass-specific anti-mouse IgG1 (1:3,000), IgG2c (1:2,000) (Invitrogen USA) or IgA (1/750) (Sigma Aldrich). As substrate 1.0 mg/ ml o-phenylendiamine (OPD, Bio Basic Canada Inc) in 0.1 M citrate-phosphate buffer, pH 5.0 containing 0.1% hydrogen peroxide was used. Optical densities (ODs) were measured with Titertek Multiskan Model 340 microplate reader (ICN, USA) at 490 nm. From the experimental protocol performed in triplicate, one representative experiment is presented in the Results.

### Avidity Assay

Avidity was measured by an ELISA elution assay as the overall strength of binding between antibody and antigen, using plates incubated for 10 min with increasing concentration of ammonium thiocyanate (NH_4_SCN) from 0 to 1 M. Antibody avidity was defined as the amount (percentage) of antibody retained for each increment of NH_4_SCN concentration.

### Ag-Specific IL-17 and IFN-γ production by Spleen Cells

After *B. pertussis* challenge, spleens from untreated and immunized mice were passed through a 40-mm cell strainer to obtain a single-cell suspension. Spleen cells (40) were seeded in 48 well culture plates in a final volume of 500 µl/well RPMI 1640 with 10% fetal bovine serum, containing 100IU/ml penicillin and 100µg/ml streptomycin. All cell samples were stimulated with heat OMV (2 µg/ml) or medium only. After 72 h of incubation (37ºC and 5% CO_2_), IFN-γ and IL-17 concentrations were quantified in supernatants by ELISA.

### Statistical analysis

The data were evaluated statistically by two-way or one-way analysis of variance (ANOVA) followed by Bonferroni for multiple comparisons (via the GraphPad Prism® software). Differences were considered significant at a p <0.05.

## Results

### Immunogenicity of OMV based vaccine prototype administered via mucosal route through homologous or heterologous vaccination scheme

To assess the immunogenicity and then the ability of OMVs to reduce *B. pertussis* colonization in the lower and upper respiratory tract, groups of mice were immunized with a 2-dose scheme administered with a formulation containing OMVs using mucosal routes (sublingual OMV_SL_ or intranasal OMV_IN_) in homologous schemes (OMV_SL-SL_ or OMV_IN-IN_) or heterologous schemes (OMV_SL-IM_, OMV_IM-SL_, OMV_IN-IM_, OMV_IM-IN_). A homologous 2-dose scheme using the intramuscular route (OMV_IM-IM_) was also incorporated for comparative analysis. For the intramuscular systemic route, we employed a previously tested and proven effective formulation comprising OMVs at a final protein concentration of 3 μg and alum as adjuvant (32,40). In the case of mucosal routes, formulations containing OMVs at a final protein concentration of 6 μg, supplemented with the mucosal adjuvant c-di-AMP, were utilized. The experimental setup and timelines of the experiment are shown in Figure 1A. High levels of serum *B. pertussis*-specific IgG were obtained with OMV_IM-IM_ and to a lesser extent in heterologous schemes, but not in homologous mucosal schemes or the non-immunized group (Figures 1BC). Intranasal immunization elicited significantly higher *B. pertussis*-specific IgG antibody responses compared to sublingual immunization (p<0.001) (Figures 1D). Comparative avidity assays conducted with different concentrations of the chaotropic agent NH_4_SCN showed that among the homologous schemes, the OMV_IM-IM_ scheme had the *B. pertussis*-specific IgG with the highest avidity (p<0.01 when 0.5M of NH_4_SCN was used) (Figure 1 EF). Increased affinity was also detected in sera obtained from the heterologous schemes, with the lowest observed in sera from mice immunized with OMV_SL-IM_ (Figure 1E).

**Figure 1.**
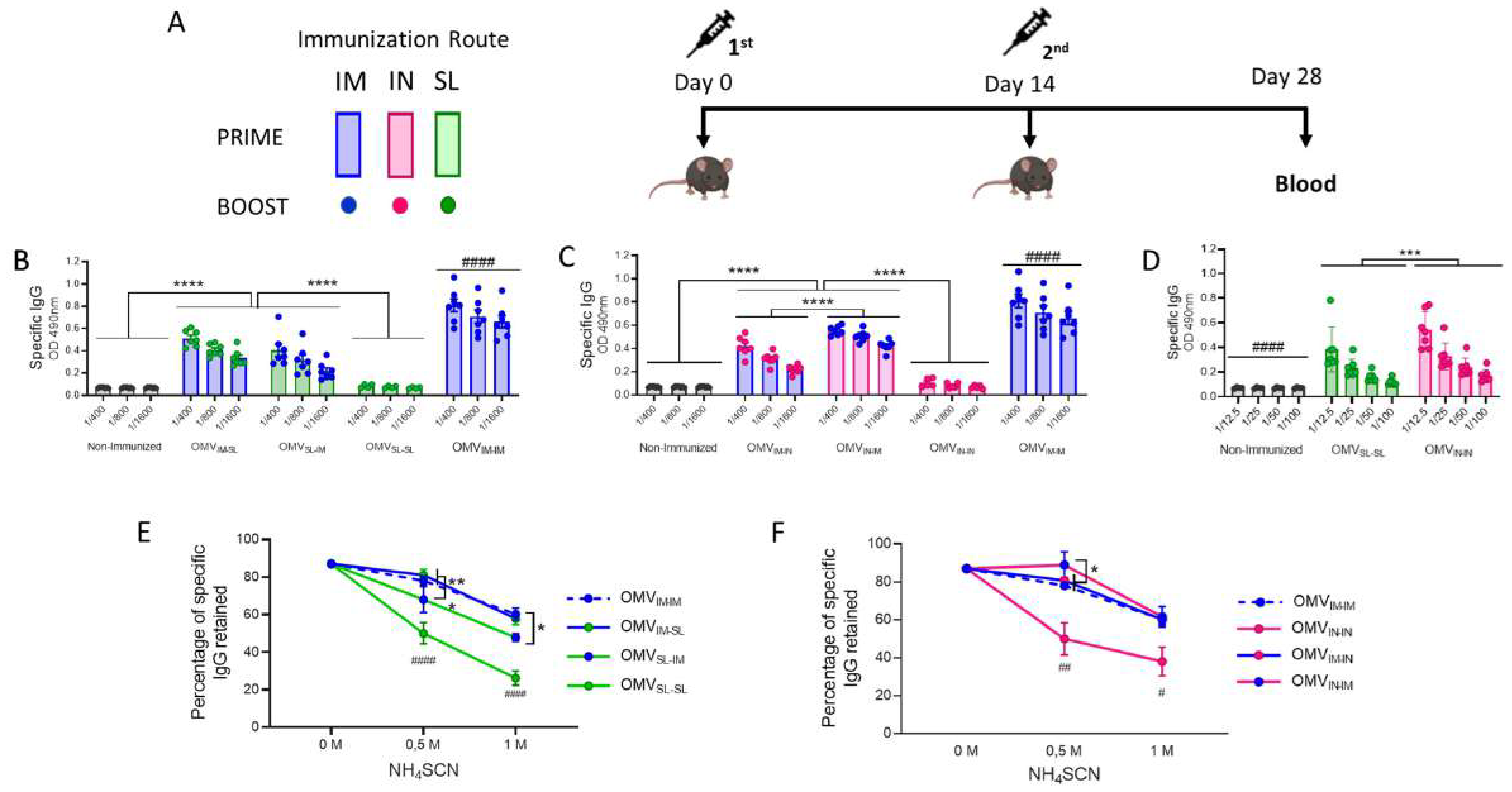
Characterization of humoral immune response induced by OMV administered through heterologous and homologous immunization schemes. **A)** C57BL/6 mice (n=7/group) were immunized on Days 0 and 14, with a 2-dose scheme administered with a formulation containing OMVs using mucosal routes (sublingual OMV_SL_ or intranasal OMV_IN_) in homologous schemes (OMV_SL-SL_ or OMV_IN-IN_) or heterologous schemes (OMV_IM-SL_ OMV_SL-IM_,, OMV_IM-IN_ or OMV_IN-IM_,). A homologous 2-dose scheme using the intramuscular route (OMV_IM-IM_) was also incorporated for comparative analysis. Different bars colors are used to discriminate the route used for the first dose and the colors of circles denote the route used for the booster dose. Specific IgG levels induced by schedules that include SL administration **(B)**, IN administration **(C)** or homologous mucosal administration **(D)** were determined in sera collected on Day 14 after the last dose (absorbance values at 490 nm for 2 sera dilutions). The avidity of IgG antibodies was also measured 14 days post the second dose and the results are presented as percentages of *B. pertussis*-specific IgG antibodies retained after exposure with 0.5 M and 1M of ammonium thiocyanate (NH_4_SCN). ^*^p<0.05, ^**^p<0.01, ^***^p<0.001, ^****^p<0.0001 by two way ANOVA using Bonferroni for multiple comparisons. The # symbol indicate significant differences between treatment and the other groups. #### p <0.0001.

Regarding the levels of *B. pertussis*-specific IgG1 (Fig 2AB), the highest levels were observed in the treatment group that received 2 doses of the OMV_IM_ formulation (Fig 2A), and in the heterologous scheme group OMV_IM-IN_ (p <0.05). The levels detected for the OMV_IM-IN_ scheme were significantly higher than those detected for OMV_IN-IM_ (p<0.0001, Fig 2B). Once again, the lowest levels of IgG1 were detected for the homologous schemes OMV_SL-SL_ or OMV_IN-IN_ and for the non-immunized control group (Fig 2AB). No difference was detected in IgG1 levels between mucosal homologous schemes (not shown).

**Figure 2.**
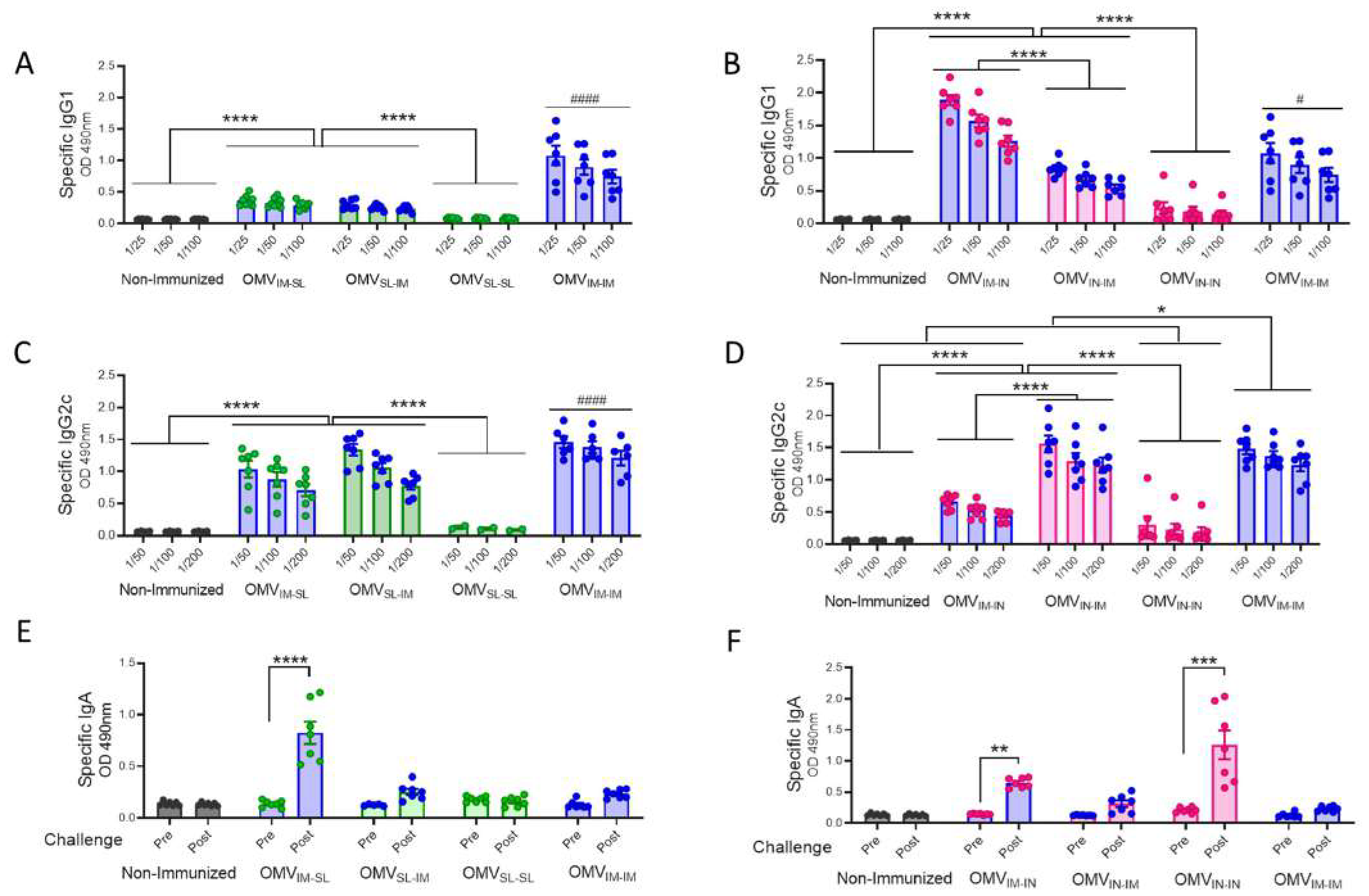
Specific *B. pertussis* IgG isotypes and IgA levels induced by OMV administered through different heterologous and homologous immunization schemes. Groups of C57BL6 mice (n=7) were immunized with a 2-dose scheme as described in panel A of Figure 1. A homologous 2-dose scheme using the intramuscular route was incorporated for comparative analysis (OMV_IM-IM_) and a group of non-immunized mice was employed as a control. Specific IgG1 **(A, B)** and IgG2a **(C, D)** levels were determined in sera of immunized mice two weeks after last dose (absorbance values at 490 nm for 3 sera dilutions). **A, C:** levels of IgG isotypes induced by schedules that include SL immunization. **B, D:** levels of IgG isotypes induced by schedules that include IN administration. IgA antibody responses in serum were evaluated before and after intranasal *B. pertussis* challenge (absorbance values at 490 nm) in the groups of immunized mice with schedules that include SL **(E)** and IN **(F)** administration. ^*^p<0.05, ^**^p<0.01, ^***^p<0.001, ^****^p<0.0001 by two way ANOVA using Bonferroni for multiple comparisons. The # symbol indicate significant differences between OMV_IM-IM_ and the other groups. # p<0.05, #### p <0.0001.

When IgG2c levels were evaluated, the highest were detected for the OMV_IN-IM_ and OMV_IM-IM_ treatments (Figure 2CD). It is worth noting, however, that the heterologous schemes including OMVSL induced significantly high levels, albeit lower than the former (Fig 2C). The IgG2c levels induced by OMV_IM-IN_ were higher than those induced by OMV_IN-IN_ (p<0.001). For mucosal homologous schemes and, as expected, the non-immunized control group, the levels of IgG2c isotypes were undetectable (Fig 2CD). An interesting finding was that for all evaluated schemes except OMV_IM-IN,_ OMV_SL-SL_ and OMV_IN-IN_, the IgG2c/IgG1 ratios were higher than 1 (supplementary material Table S1), suggesting the induction of a Th1-directed immune response.

IgA antibody responses were only detected following intranasal *B. pertussis* challenge in mice treated with homologous intranasal immunization scheme and in those receiving the booster dose via mucosal route (OMV_IM-IN_ and OMV_IM-SL_ heterologous treatments), while they were barely detectable in the other tested groups (Fig 2 EF).

IFN-γ (Th1 profile marker), and IL-17 (Th17 profile marker) levels were also evaluated through splenocytes stimulation assays in mice challenged with a *B. pertussis* suspension (Figure 3A). The highest levels of OMV-specific IFN-γ and IL-17 were detected when OMV_IN_ was included, either in a homologous or heterologous routes of immunization, with no significant differences observed between the treatments that included OMV_IN_ (Fig 3BC). It is noteworthy that the levels of IL-17 detected in the schemes where OMV_IN_ is present are significantly higher than those of non-immunized challenged mice (p<0.001). It was also found that IL-17 levels in the heterologous scheme OMV_SL-IM_ were significantly higher than those induced by the other treatments containing any dose of OMV_SL_, as well as compared to non-immunized challenged animals (p<0.001).

**Figure 3.**
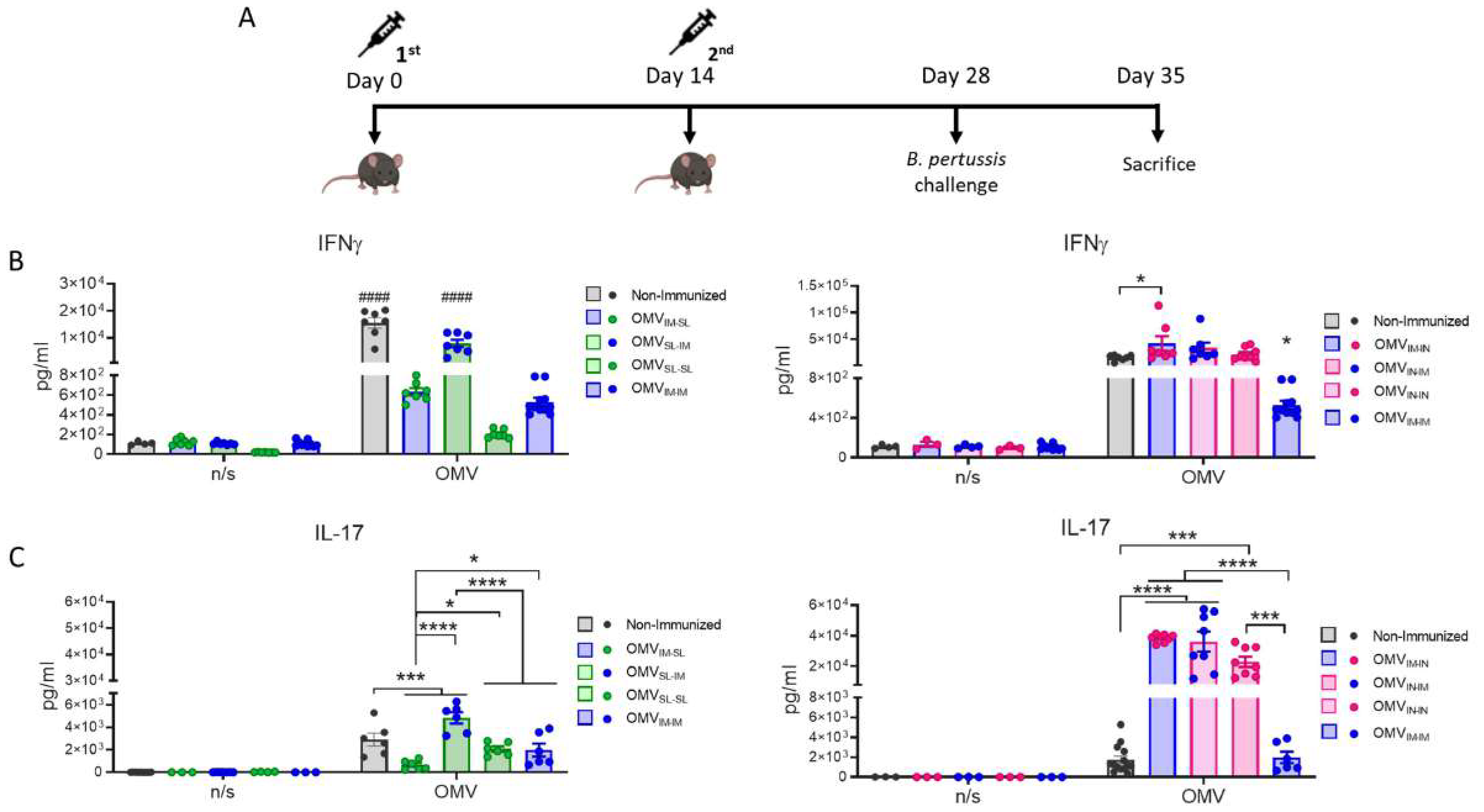
Characterization of cellular immune response induced by OMV administered through heterologous and homologous immunization schemes. **A)** Mice were immunized on Day 0 and 14, with a 2-dose scheme administered with a formulation containing OMVs using mucosal routes in homologous schemes or heterologous schemes, as described in Figure 1. Seven days after challenge with a sublethal dose (1×107-5×107 CFU/40ul), mice were sacrificed, and their spleen cells were restimulated *ex-vivo* with 2μg/ml of OMV derived from *B. pertussis* or non-stimulated as controls (n/s). Levels of secreted IFN-γ **(B)** and IL-7 **(C)** following splenocytes stimulation with medium or OMVs were determined by ELISA. Bars are means ± SEM of pg/ml. ^*^p<0.01, ^***^p<0.001, ^****^p<0.0001 by two way ANOVA using Bonferroni for multiple comparisons. The # symbol indicate significant differences between the treatment and the other groups. #### p <0.0001.

All these results demonstrate that the tested mucosal immunization schedules induced a robust specific humoral immune response when they were used in heterologous schemes showing higher levels of specific IgG in comparison with those detected for homologous mucosal schemes. It was also observed that the OMVs used in homologous or heterologous mucosal administration schemes induce a mixed Th1/Th17 profile, although more pronounced in schemes containing OMV_IN_ and also in the heterologous scheme OMV_SL-IM_. High levels of IgA were detected for OMV_IN-IN_ and for heterologous schemes that include mucosal administration for the second dose (OMV_IM-SL_ and OMV_IM-IN_).

### Protective capacity of OMV based vaccine prototype administered via mucosal route through homologous or heterologous vaccination scheme

To assess the protective capacity exhibited by the different vaccination strategies tested here, groups of animals receiving 2-dose regimens were intranasally challenged with a suspension of *B. pertussis* at sublethal dose (1×107 – 5×107 CFU/40 μl) 14 days after receiving the last dose (Figure 3A). Seven days post-challenge, the mice, including those in the non-immunized group, were euthanized by cervical dislocation to evaluate bacterial colonization in the upper and lower respiratory tract. As expected, the non-immunized control group had a highest level of bacteria: 4.91×10^4^ CFU/lungs and 4.48×10^4^ CFU/nose (Fig 4 AB). The majority of immunized mice showed a significant reduction in the bacterial load in the lungs compared to non-immunized animals (p<0.0001), with the smallest reduction observed in the OMV_SL-SL_ immunized mice (reduction of 1.25 logs). For the heterologous treatments OMV_SL-IM_ and OMV_IM-SL_, significant reductions of 2.58 and 2.37 logs, respectively, were recorded, and in the case of treatments that include a mucosal dose intranasally, reductions were 1.67 for mice immunized with both OMV_IM-IN_ and OMV_IN-IM_ (Figure 4B). It is noteworthy that in the homologous schemes OMV_IN-IN_ and OMV_IM-IM_, reductions were at least 3 logs, similar to what was observed for commercial aP (reduction of 2.99 logs, Figure 4C) and wP (reduction of 3.25 logs, Figure 4F) vaccines administered intramuscularly.

**Figure 4.**
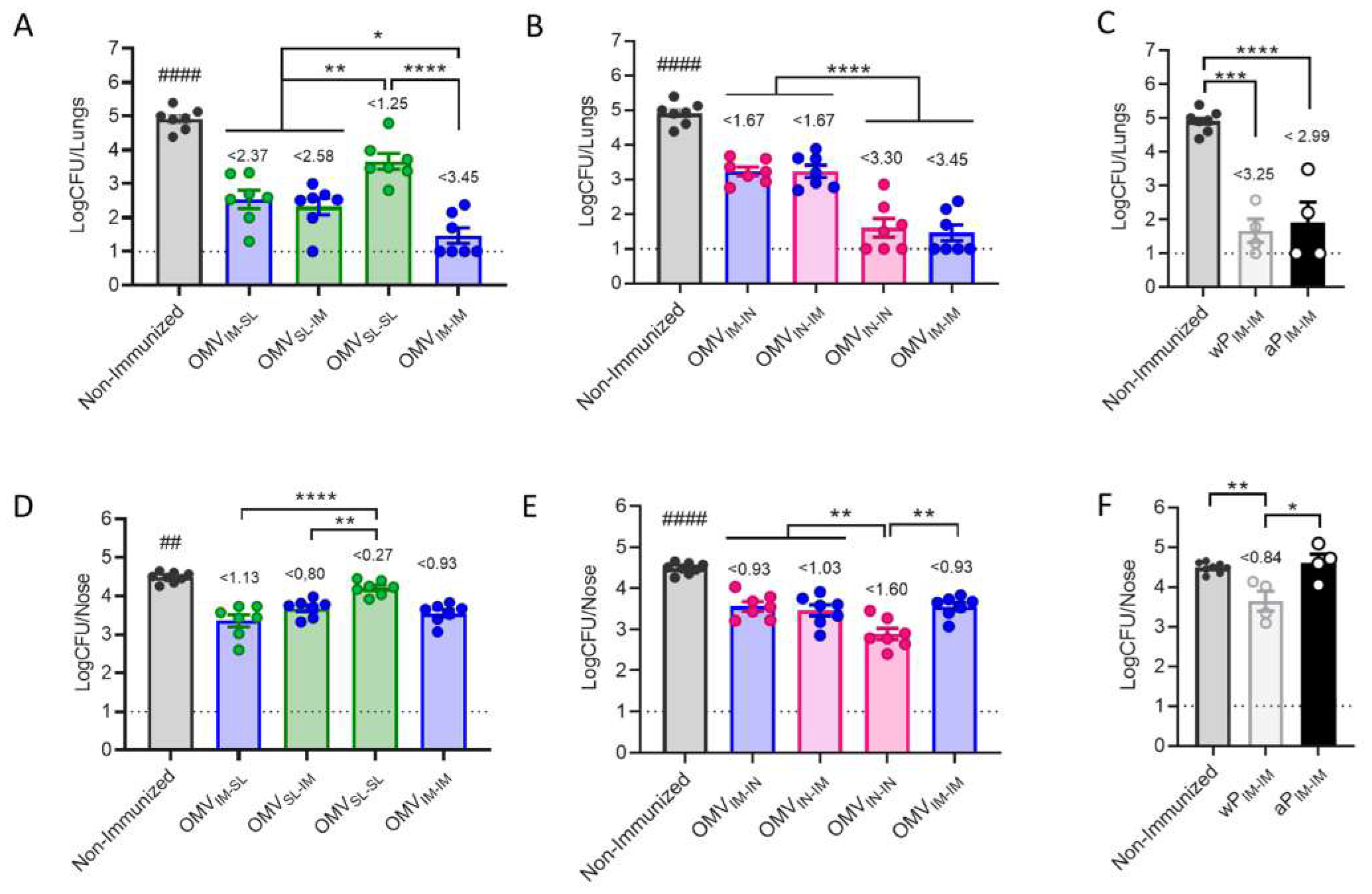
Effect of OMV mucosal homologous and heterologous immunization on protection against *B. pertussis* infection. C57BL/6 mice (n=7/group) were immunized on Days 0 and 14, with a 2-dose scheme OMV administered as described in panel A of Figure 1. Groups of animals immunized with commercial aP, or commercial wP vaccines (n=8 in each group) or non-immunized were included as positive and negative protection control. Mice from all groups were challenged with a sublethal dose (1×10^7 – 5 x10^7 40 μl-1) *B. pertussis* Tohama phase I 14 days after the second dose followed by sacrifice 7 days after challenge. The number of bacteria recovered from mouse lungs **(A, B and C)** or nose **(D, E and F)** expressed as the log10 of CFUs per lungs or nose, is plotted on the ordinate, the different treatments here tested are indicated on the abscissa, with the data representing the means ± the SD. The dotted horizontal line indicates the lower limit of detection. The reduction detected in protection levels induced by different formulations in comparison with non-immunized animals is indicated at the top of the figures. ^*^p<0.05, ^**^p<0.01, ^***^p<0.001 ^****^p<0.0001 by one way ANOVA using Bonforroni for multiple comparisons.. The # symbol indicate significant differences between the treatment and all the other tested groups. ## p<0.01, #### p <0.0001.

Similar results were observed regarding the ability to reduce bacterial colonization in the upper respiratory tract (nose) (Figure 4DE). The heterologous treatments showed a reduction of at least 0.8 logs (Figure 4 DE), similar to that detected for mice that received 2 doses of wP_IM_ (reduction of 0.84 logs) (Figure 4 F). The heterologous scheme that induced the highest reduction was OMV_IM-SL_ with a reduction of 1.13 logs. While the homologous schemes OMV_IN-IN_ (reduction of 1.60 logs) and OMV_IM-IM_ (reduction of 0.93 logs) reduced bacterial colonization in the nose more effectively than the commercial aP vaccine applied in a 2-dose intramuscular scheme (non reduction) (p<0.001), OMV_SL-SL_ did not (Figure 4D).

In summary, while homologous immunization with OMV_IN-IN_ showed the highest protective capacity in both upper and lower airways, heterologous schemes, particularly OMV_IM-SL_, exhibited adequate levels of protection surpassing the protection conferred in the nose by the commercial cellular vaccine.

### Immunogenicity and protective capacity of intranasal homologous scheme using formulation containing different adjuvants

With the aim of assessing whether the combination of adjuvants c-di-AMP with alum can improve the efficacy of the best scheme identified thus far in this study, OMV_IN-IN_, we conducted comparative assays using formulations containing the adjuvant combination versus those with a single adjuvant. By these assays we detected that *B. pertussis* IgG levels induced by the OMVs formulated with alum+c-di AMP were the lowest though they did differ from those detected in non-immunized mice (Figure 5A). It was also observed that the antibody affinity was lower in sera from mice intranasally administered with 2 doses of OMVs formulated with combined adjuvant or alum compared to those detected for c-di AMP formulation (Figure 4B).

**Figure 5.**
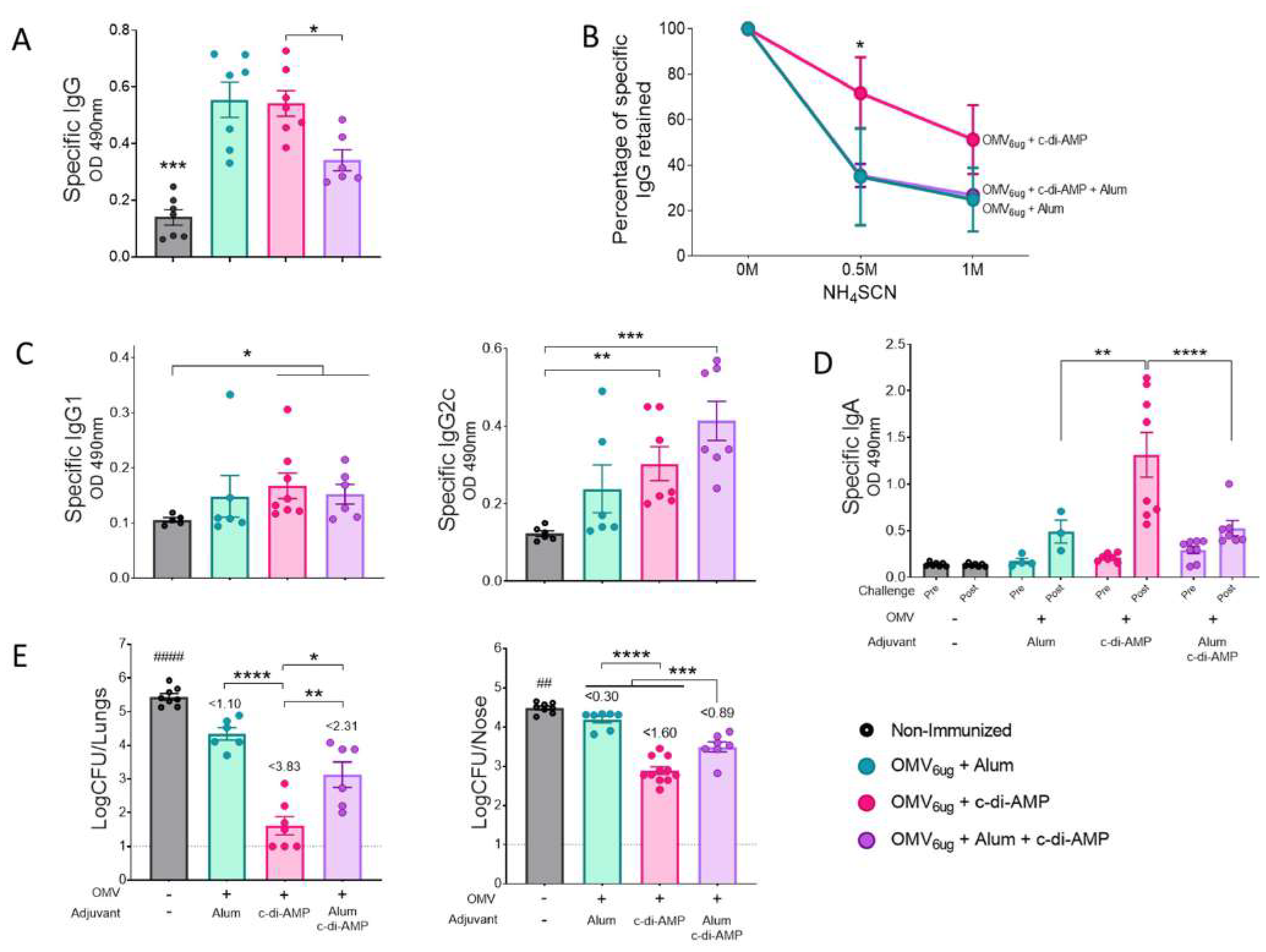
Characterization of immunogenicity and the protection capacity induce by OMVs administered by intranasal route using different adjuvants. Groups of C57BL6 mice (n=7) were immunized with a 2-dose scheme administered IN with a formulation containing OMVs formulated with c-di AMP, Alum, or c-di AMP plus Alum. As a control, a group of non-immunized were employed. **A**. Specific IgG levels induced by different schedules were determined in sera collected on Day 14 after the last dose (absorbance values at 490 nm). The avidity of IgG antibodies was also measured 14 days after the second dose and the results are presented as percentages of *B. pertussis*-specific IgG antibodies retained after exposure with 0.5 M and 1M of ammonium thiocyanate (NH_4_SCN). **B**. Specific IgG1 and IgG2a levels were determined in sera of immunized mice two weeks after last dose (absorbance values at 490 nm). Mice from all groups were challenged with a sublethal dose (1×10^7 – 5 x10^7 40 μl-1) *B. pertussis* Tohama phase I 14 days after the second dose followed by sacrifice 7 days after challenge. **C**. IgA antibody responses in serum were detected before and after intranasal *B. pertussis* challenge (absorbance values at 490 nm). **D**. The number of bacteria recovered from mouse lungs (left panel) or nose (right panel) expressed as the log10 of CFUs per lungs or nose, is plotted on the ordinate, the different treatments here tested are indicated on the abscissa, with the data representing the means ± the SD. The reduction detected in protection levels induced by different formulations in comparison with non-immunized animals is indicated at the top of the figures. The dotted horizontal line indicates the lower limit of detection. p<0.05, ^**^p<0.01, ^***^p<0.0001, ^****^p<0.00001 by one way ANOVA using Bonforroni for multiple comparisons.

Mice immunized with formulations containing c-di AMP and c-di AMP+alum as adjuvants also exhibited *B. pertussis* IgG1 and IgG2c levels that differed slightly from those of non-immunized animals (see Figure 5C). The former scheme also showed the highest increase in IgA levels (Figure 5D). Regarding the IgA increase, once again, the groups of mice immunized with the formulation containing alum (either alone or in combination with c-di AMP) presented the lowest value (see Figure 5D). Regarding the ability of these formulations to reduce bacterial colonization in the upper and lower respiratory tracts after 7 days of challenging mice with a sublethal concentration of *B. pertussis* (Figure 5 E), we observed that mice immunized with the formulation containing only c-di AMP presented the highest reduction. In the lower respiratory tract, the reduction in bacterial colonization for this treatment compared to non-immunized animals was 3.83 logs, and in the upper respiratory tract, the reduction was 1.60 logs. The formulation containing the mixture of adjuvants also induced a significant reduction in bacterial colonization in both the lower and upper respiratory tracts, with reductions of 2.31 logs and 0.89 logs, respectively. The lowest reduction in the upper respiratory tract was observed in animals treated with OMV formulated with alum (Figure 5 E)

## Discussion

We have previously demonstrated that *B. pertussis* OMVs, comprising a significant array of immunogens and PAMPs, constitute a safe and effective vaccine prototype for preventing bacterial colonization in the lower respiratory tract when administered either parenterally or intranasally (32,41). In a recently submitted manuscript, we also demonstrated the adjuvant effect of OMVs when given via both systemic and mucosal routes. In this study, we assessed the efficacy of OMVs derived from *B. pertussis* when administered via mucosal immunization (sublingual or intranasal routes) to confer protective immunity against colonization, not only in the lower respiratory tract but also in the upper respiratory tract of mice. We examined both homologous and heterologous 2-dose schedules. The intramuscular route was included in heterologous regimens, and its inclusion in homologous regimens was for comparative purposes.

While many questions regarding sublingual immunization still need to be addressed, including those related to the use of suitable adjuvants and the optimization of vaccine formulation to further enhance efficacy, it has several precedents in animal models demonstrating its safety and effectiveness against various respiratory pathogens. (influenza, RSV, SARS-CoV-2) (42,43). Concerning intranasal vaccination, the understanding that the nasal cavity harbors abundant lymphatic tissue, which triggers both humoral responses— where immunoglobulin A comprises over 15% of total immunoglobulins—and cellular immune responses, has led to numerous studies verifying its safety and efficacy (44). Here, we have found that OMVs formulated with alum and administered via intramuscular injection, or formulated with c-di-AMP and administered via the sublingual mucosal route in a heterologous administration scheme, or intranasally in both homologous and heterologous schemes, effectively reduce bacterial colonization in both the lower respiratory tract (associated with severe disease) and the upper respiratory tract (implicated in transmission). Similar to commercial pertussis vaccines wP and aP currently used in various countries, schemes that included homologous administration of two doses of OMVs administered via IM or IN achieved a reduction in *B. pertussis* lung colonization of approximately 3 logs compared to the CFU levels detected in non-immunized animals. An important finding is that these schemes, unlike aP vaccines that failed to reduce bacterial colonization in the nose, achieved a reduction of almost 1 log in the case of the OMV_IM-IM_ scheme and 1.60 logs for the OMV_IN-IN_ scheme. Regarding the reduction of bacterial colonization induced by the commercial wP vaccine, while the reduction in colonization was similar to that of the OMV_IM-IM_ scheme, the reduction induced by the OMV_IN-IN_ scheme was significantly superior (p<0.001), positioning this scheme above those of the commercially available vaccines currently in use. This result regarding intranasal vaccination aligns with findings reported by Raeven et al (34), who conducted comparative studies between vaccination schemes using OMVs derived from another strain of *B. pertussis* administered subcutaneously and intranasally. The colonization levels achieved by these authors, however, differed from those reported here, likely due to the use of a different strain of *B. pertussis* and variations in the challenge doses (34).

Furthermore, here for the first time we detected that heterologous schemes such as OMV_IN-IM_, OMV_IM-IN_, and OMV_IM-SL_ are also capable of reducing bacterial colonization by at least 1.67 logs in the lower respiratory tract and at least 0.93 logs in the upper respiratory tract. The OMV_SL-IM_ scheme also provided protection against lung colonization, reducing levels compared to the non-immune mouse group by 2.58 logs in the lungs and 0.80 logs in the nose, slightly less than the counterpart OMV_IM-SL_ scheme. The least immunogenic and protective scheme turned out to be the homologous OMV_SL-SL_ scheme. Heterologous schemes that included doses administered via SL, although immunogenic, showed a lower capacity to induce humoral or cellular responses compared to homologous or heterologous schemes that included doses administered intranasally. Interesting, we observed that the heterologous schemes where the second dose was administered by intramuscular route, the induced IgG2c levels were significantly higher than those detected in any of the schemes starting with an intramuscular dose. For these schedules the IgG2c/IgG1 ratio was higher than 1 (supplementary material Table S1). On the other hand, heterologous schemes that utilize the mucosal route for administering the second dose, along with the homologous OMV_IN-IN_ scheme, induce elevated levels of IgA post-challenge. Moreover, schemes incorporating the intranasal route for the second dose (whether homologous or heterologous) elicit high levels of IL-17, which, together with IgA, are described as crucial for inducing protection in the upper respiratory tract. (45). Solans et al. (45) reported results indicating significant differences in the protective mechanisms between the upper and lower murine respiratory tract, highlighting the IL-17-dependent and IgA-mediated mechanism for protecting against *B. pertussis* nasopharyngeal colonization. Schemes in which the second dose is administered via the mucosal route have also been reported as protective for other pathogens (46,47).

In the context of intranasal immunization, we identified that the OMV vaccine formulated with c-di AMP as adjuvant promoted a superior humoral response (IgG and IgA post-challenge) compared to that detected in animals treated with OMV formulated with c-di-AMP+Alum. The specific IgG induced by OMV +c-di AMP also exhibited a higher affinity than those generated by the one containing c-di AMP+Alum or alum alone.

Consistent with published results using the baboon model, our findings in the mouse model provide additional evidence of the limitations of aP vaccines formulated with alum in reducing bacterial colonization in the nasal cavity (27) (31). While immunization with an alum-adjuvanted aP vaccine administered parenterally (IM) effectively prevented lung infection, it did not prevent nose colonization by *B. pertussis*. Moreover, when the aP vaccine is administered intranasally, protection against colonization in both lungs and nose is highly deficient; the colonization levels do not differ from those in the non-immunized group (supplementary material Figure S1). In contrast, the OMV formulated with c-di AMP conferred higher immunity against nasal colonization while maintaining excellent protection against lung infection. The least effective formulation administered intranasally against both lung and nose colonization was the one containing OMV and alum.

In summary, our findings suggest that intranasal immunization with OMV vaccine prototype formulated with c-di AMP in both homologous and, to a lesser extent, heterologous administration schedules, is an effective approach for inducing humoral, IL-17, and IFN-γ responses, thereby providing protection against nasal and lung colonization by *B. pertussis*. The presented results offer proof-of-principle for the use of OMV, whether administered intramuscularly or intranasally, surpassing current vaccines in use, particularly the acellular vaccine. This supports the foundation of a third-generation pertussis vaccine for humans.

## Legends to the Figures

**SUPPLEMENTARY TABLE 1**. IgG2c and IgG1 levels and IgG2c/IgG1 ratio for homologous and heterologous immunized mice.

**SUPPLEMENTARY FIGURE 1. Effect of aP immunization on protection against *B. pertussis* infection**. C57BL/6 mice were immunized on Days 0 and 14, with a 2-dose scheme aP administered by intramuscular or intranasal route. Mice from all groups were challenged with a sublethal dose (1×10^7 – 5 x10^7 40 μl-1) *B. pertussis* Tohama phase I 14 days after the second dose followed by sacrifice 7 days after challenge. The number of bacteria recovered from mouse lungs **(A)** or nose **(B)** expressed as the log10 of CFUs per lungs or nose, is plotted on the ordinate, the different treatments here tested are indicated on the abscissa, with the data representing the means ± the SD. The dotted horizontal line indicates the lower limit of detection. The reduction detected in protection levels induced by different formulations in comparison with non-immunized animals is indicated at the top of the figures. ^****^p<0.0001 by one way ANOVA using Bonforroni for multiple comparisons.

