## Supplemental Table 1 for "Use of Mucosally Administered Outer Membrane Vesicles Derived from *Bordetella pertussis* to Diminish Nasal Bacterial Colonization"

**SUPPLEMENTARY TABLE 1.** IgG2c and IgG1 levels and IgG2c/IgG1 ratio for homologous and heterologous immunized mice.

| | **$OMV_{IM-IM}$** | **$OMV_{SL-SL}$** | **$OMV_{IM-SL}$** | **$OMV_{SL-IM}$** | **$OMV_{IN-IN}$** | **$OMV_{IM-IN}$** | **$OMV_{IN-IM}$** |
|---|---|---|---|---|---|---|---|
| **IgG2c** | 1,460 | ND | 1.039 | 1.342 | ND | 0.661 | 1.563 |
| **IgG1** | 0.894 | ND | 0.327 | 0.255 | ND | 1.570 | 0.679 |
| **IgG2c/IgG1** | 1.63 | - | 3.18 | 5.27 | - | 0.42 | 2.30 |
