## Supplementary figures and images for "Use of Mucosally Administered Outer Membrane Vesicles Derived from *Bordetella pertussis* to Diminish Nasal Bacterial Colonization"

### supplemental Fig 1

Supplementary Figure S1

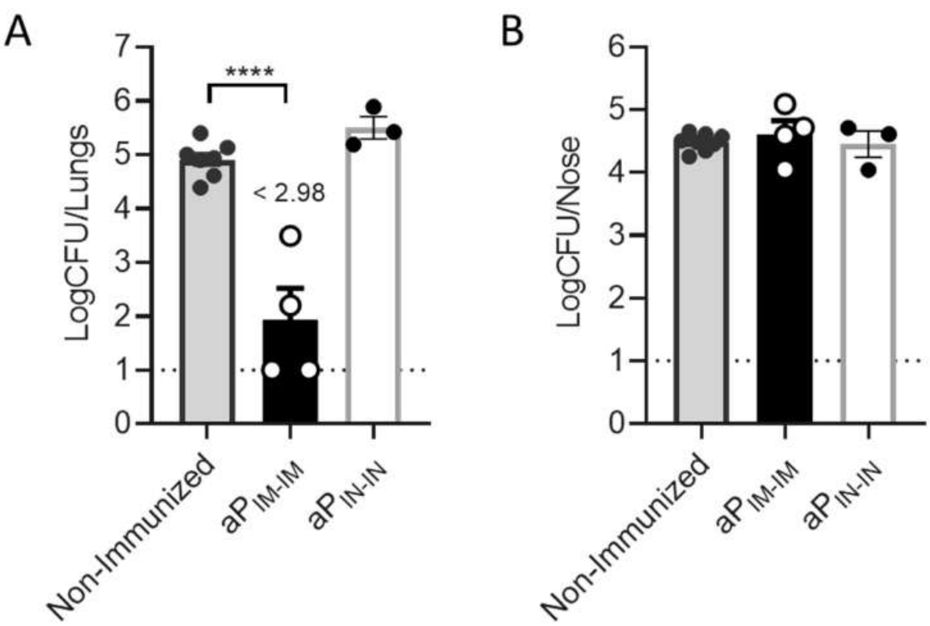
